# Fine-scale niche partitioning of prokaryotic communities across the deep chlorophyll maximum coenocline

**DOI:** 10.64898/2026.08.11.744127

**Authors:** Marta Sebastián, Carolina Marín-Vindas, Aleix Obiol, Clara Cardelús, Vanessa Balagué, Isabel Ferrera, Olga Sánchez, Josep M Gasol

## Abstract

The Deep Chlorophyll Maximum (DCM) is likely the most important feature organizing the marine epipelagic environment. Within this layer, opposing gradients of light and nutrients create a stratified habitat that supports high phytoplankton biomass and a substantial fraction of oceanic primary production. Despite its ecological importance, most studies treat the DCM as a single depth, overlooking its fine-scale heterogeneity. Here we investigated prokaryotic community organization across the DCM in the northwestern Mediterranean Sea through high-resolution sampling of four profiles collected over two days. Free-living (0.2-3 µm) and particle-associated (3-20 µm) communities were characterized using 16S rRNA gene amplicon sequencing. Prokaryotic communities changed progressively along the vertical gradient, revealing the DCM as a microbial coenocline with continuous community turnover. Fuzzy clustering identified distinct assemblages associated with environmental transitions from warm surface waters to the chlorophyll maximum, the nitrite peak below the DCM, and deeper nitrate-rich layers. In both the free-living and particle-associated fractions, most ASVs remained consistently associated with the same depth-defined clusters across all samplings, indicating stable niche partitioning over short timescales. However, these temporally stable ASVs accounted for a substantially smaller fraction of community sequences in particle-associated communities, suggesting higher dynamism, likely driven by particle-mediated transport. Nevertheless, phylogenetic analyses revealed that closely related ASVs tended to occupy similar depth niches, indicating that habitat preferences are phylogenetically conserved in both size fractions. Our results demonstrate prokaryotic niche partitioning over scales of only a few meters within the DCM, highlighting the importance of fine-scale sampling for understanding microbial community structure and responses to ocean change.

## Introduction

The Deep Chlorophyll Maximum (DCM) is a subsurface layer with maximum chlorophyll *a* concentrations, typically found in stratified oceans. It develops at depths where light levels remain sufficient for photosynthesis while nutrients supplied from deeper waters, through diffusion or upwelling, sustain phytoplankton growth (Cullen, 1982; Estrada et al., 1993; Latasa et al., 2017). Because its formation requires a stable water column, the DCM is a characteristic feature of tropical and subtropical regions, where it can remain for long periods of time. In contrast, in temperate oceans, it is generally seasonal and appears after water column stratification occurs (Cornec et al., 2021). The DCM can support a substantial proportion of the water column’s primary production, accounting for up to 50-60% in highly stratified waters (Lorenzo et al., 2004; Weston et al., 2005). Together with its proximity to export depth horizons (Guidi et al., 2016; Tilstone et al., 2017), this makes the DCM a major contributor to the biological carbon pump.

DCM-centered studies have largely focused on the mechanisms governing their formation and persistence (Cornec et al., 2021; Cullen, 1982; Moeller et al., 2019), and the vertical distribution of major phytoplankton groups using pigment-markers, flow cytometry or fluorescent *in situ* hybridization (Cabello et al., 2016; Latasa et al., 2017, 1992; Zubkov et al., 2000). In contrast, other microbial community members, such as heterotrophic prokaryotes and non-pigmented eukaryotes, have often been examined by considering the DCM as a single sampling depth (Beman et al., 2011; Giner et al., 2020; Medina-Silva et al., 2018; Mestre et al., 2018; Sunagawa et al., 2015; Walsh et al., 2015). Despite this limited vertical resolution, diversity surveys have consistently shown that DCM assemblages differ markedly from those of surface waters. However, by reducing the DCM to a single sampling point, these studies overlook the pronounced environmental gradients that characterize the DCM layer, and fail to capture its full spatial heterogeneity. Indeed, the DCM could function as a coenocline (Whittaker, 1960): a dynamic ecological transition zone between well-lit surface waters and the underlying light-limited environment, integrating varying features of both biomes.

Consistent with this view, the limited number of high-resolution DCM studies conducted so far have revealed strong vertical structuring of microbial communities within the DCM, including both prokaryotes (Gazulla et al., 2026; Haro-Moreno et al., 2018) and eukaryotes (Cabello et al., 2016; Dolan and Marrasé, 1995; Latasa et al., 2017). For example, a detailed metagenomic survey at a single location in the western Mediterranean based on samples collected at 15 m depth intervals across the stratified photic layer, revealed pronounced changes in the free-living prokaryotic community composition over relatively short vertical distances (Haro-Moreno et al., 2018). However, because this study performed a single sampling during the stratification period, it could not evaluate whether these vertical patterns remain stable over short timescales. Such temporal variability is plausible given that the processes underlying DCM formation and maintenance, including phytoplankton photoacclimation (Becker et al., 2021; Steele, 1962), or nutrient availability and grazing pressure (Moeller et al., 2019), fluctuate over the diel cycle in response to changes in irradiance and biologically driven circadian rhythms. Picophytoplankton exhibit depth-dependent diel division patterns throughout the photic zone, including the DCM (Vaulot et al., 1995), and bacterial activity similarly follows diel oscillations (Gasol et al., 1998; Ruiz-González et al., 2012). Together, these observations suggest that the vertical organization of microbial communities within the DCM may itself undergo diel shifts. Moreover, the pronounced stratification patterns observed in free-living prokaryotic communities may not apply to particle-associated prokaryotes which can be governed by surface-determined ecological factors (Ruiz-González et al., 2020). These two fractions represent distinct ecological lifestyles, differing in taxonomic composition, functional capabilities, and the environmental factors shaping their distribution (Roth Rosenberg et al., 2021). As a result, particle-associated prokaryotes could exhibit their own distinct spatial and temporal dynamics across the DCM.

Here, we investigated the fine-scale vertical organization of prokaryotic communities across the DCM in the northwestern Mediterranean Sea through high-resolution vertical sampling, with twelve samplings depths spanning the DCM structure. Samples were collected at a single station over two consecutive days, at 10 h and 20 h GMT, to evaluate whether the pronounced vertical gradients previously observed in stratified waters remain stable over short timescales. We analyzed prokaryotic community composition using amplicon sequence variants (ASVs) in both the free-living (0.2-3.0 µm) and particle-associated (3.0-20 µm) fractions. Specifically, our objectives were i) to characterize the fine-scale distribution of prokaryotic taxa across the DCM, ii) to compare the vertical patterns between free-living and particle-associated communities, and iii) to determine whether these distributions exhibit short-term temporal variability within the highly dynamic DCM feature.

## Materials and methods

### Study area and sample collection

Sampling was performed during the REMEI cruise on board the *R/V* García del Cid from September 27^th^ to September 29^th^, 2017 centered around 40° 48.99 N 3° 4.93 E, in the northwestern Mediterranean Sea between the Barcelona coast and the island of Mallorca. Because the research vessel was not equipped with a dynamic positioning system, slight drift occurred during sampling. Consequently, the actual sampling locations differed slightly from the intended station coordinates, with all four samplings located within a 500 m radius. Four vertical casts were performed with a CTD profiler equipped with a SBE 43 dissolved oxygen sensor and a Seapoint fluorometer. Sampling took place on September 27^th^ at ca. 20h, on September 28^th^ at ca. 10h, and on September 29^th^ at 10h and 20h (GMT). Irradiance was measured with a PUV-2500 (Biospherical Instruments) radiometer at noon. Water samples were collected using a 12-bottle Niskin rosette. High-resolution sampling was conducted with samples collected every 5-10 m within the DCM layer. Additional samples were obtained from surface waters and from depths below the DCM (around 160-175 m) during each cast. To minimize disturbance of the stratified physical and biological structure of the water column, the rosette was retrieved at a low ascent rate, and the Niskin bottles were triggered sequentially without stopping the rosette retrieval, as in Cabello et al., (2016). Details on methods related to Chlorophyll a, inorganic nutrient measurements, heterotrophic bacteria and picophytoplankton abundance, DNA extraction and 16s amplicon sequencing and processing can be found in the Supplementary information.

### Data analyses

Data treatment was performed using R (4.4 version). For diversity analyses the samples were randomly subsampled to the lowest number of reads (4.795 and 5.694 reads for the 0.2-3 µm and 3-20 µm size fractions, respectively) using the *rarefy* function in the *vegan* R package (Oksanen et al., 2019). Prokaryotic richness (the number of ASV per sample) and sample evenness (using the Pielou index (*J* = *H*/ln(nASV), where H is the Shannon index and nASV is the richness in every sample), were calculated using the *vegan* package (Oksanen et al., 2019). Non-metric Multidimensional Scaling (NMDS) was performed with the *vegan* package using Bray-Curtis distances of rarefied ASV abundance tables.

The vertical spatial patterns of the communities were analyzed by Fuzzy C-Means Clustering (Bezdek, 1981), which, for each ASV, assigns a membership score for all clusters concurrently (from 0 to 1). Fuzzy clustering was performed individually for each DCM and size fraction combination using the *cmeans* function (*e1071* package, v. 1.7.9; (Meyer et al., 2021)). The fuzzifier parameter *m* was estimated following Schwämmle and Jensen (2010), resulting in a value of 1.2797 ± 0.018 (mean ± sd). To choose the number of clusters (k), we tested several cluster selection indices: Within Cluster Sum of Squared Error, Simple Structure Index (*cascadeKM* function, *vegan,* Dimitriadou et al., (2002)) and Normalized Partition Coefficient (*vegclustIndex* function, *vegan,* Bezdek, (1981)), yet these indices provided very contrasting results (from 2 to 10 clusters). According to Bagnaro et al. (2020) and due to mathematical limitations, the number of fuzzy clusters cannot exceed n/2 − 1, where n is the total number of observations. Since we had 12 observations, the maximum number of clusters would be 5. We examined the vertical distribution patterns of 2, 3, 4, and 5 clusters and visually evaluated the results. The final number of clusters (5 for the 0.2 – 3 µm size fraction and 4 for the 3 -20 µm size fraction) was selected based on clear distinct trends, and choosing the number of clusters that yielded a higher proportion of ASVs with a high membership score, while avoiding redundant clusters (Fig. S1). For this analysis, only ASVs representing at least 0.1% of reads in one sample were considered. To group ASVs based on their spatial distribution while minimizing the influence of abundance levels, we applied a Centered Log-Ratio (CLR) transformation to their abundances before the clustering and each ASV’s transformed abundance was then centered and scaled by subtracting its mean and dividing by its standard deviation. To investigate associations between the different clusters and environmental variables, we performed Mantel tests between the summed relative abundance of the ASVs within each cluster and the environmental variables, using the *vegan* package.

### Phylogenetic signal in the vertical distribution of ASVs

We calculated the phylogenetic distance among ASVs to determine whether their positions across the DCM structure exhibited a phylogenetic signal. To this end, ASV sequences were aligned with the *mafft* software and phylogenetic relationships were inferred with iq-tree (Minh et al., 2020), using the model GTR+G+FO. The generated tree file was read in R with the *ape v5.8-1* package (Paradis and Schliep, 2019) and distances extracted with the function *cophenetic.* The optimal position of each ASV across the DCM was defined as the abundance-weighted mean depth derived from its vertical distribution profile. Plots were generated with *ggplot2 (Wickham, 2016)*.

## Results

### Environmental transitions along the DCMs

Despite some variability in the depth and shape of the fluorescence maximum, the four vertical profiles showed a consistent overall structure, characterized by a broad DCM extending between approximately 50 and 80 m depth. During the daytime samplings, the 1% photosynthetically active radiation (PAR) level was located at ∼70 m, within the DCM, while the 0.5% and 0.1% PAR levels were reached at approximately 90 and 100 m, respectively (data not shown). Fluorescence and oxygen concentration followed a similar pattern, increasing below the upper mixed layer (Fig 1, Fig. S2) and slowly decreasing between 80 and 100 m depth (Fig. 1). The first profile (DCM1, Fig. 1) showed a fairly broad DCM, with two measured similar chlorophyll peaks at 60 m (0.371 mg m^-3^) and 85 m (0.367 mg m^-3^, Fig. 2, Table S1). In the second profile the DCM was more defined (DCM2, Fig. 1), showing maximum values of Chl a at 90 m (0.380 mg m^-3^, Fig. 2, Table S1). In the third sampling the fluorescence profile revealed a broader DCM (DCM3, Fig. 1), with the highest chlorophyll concentration recorded at 56 m (0.357 mg m^-3^, Fig. 2, Table S1). This peak roughly coincided with a maximum in oxygen concentration and a minimum in salinity (DCM3, Fig. 1 and Fig. S2. The fourth profile showed the narrowest DCM, with the highest chlorophyll concentrations measured between 45 and 82 m, with a peak at 58 m (0.345 mg m^-3^, Fig. 2, Table S1).

**Figure. 1.**
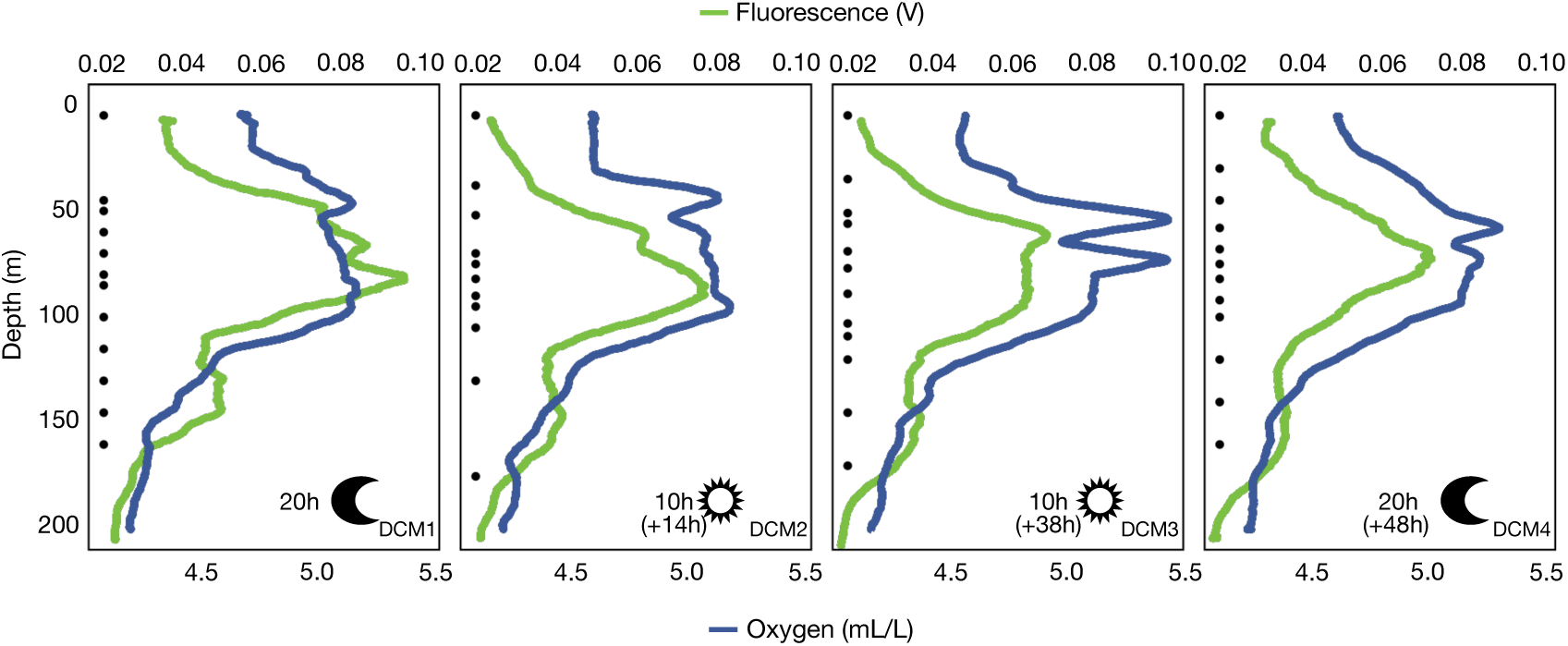
Vertical profiles of fluorescence and oxygen across the four DCM samplings performed at a single station in the northwestern Mediterranean Sea. The sampling periods (day/night), sampling hour and the time lag since the first sampling (in parenthesis) are shown for each profile. Black dots indicate the depths at which seawater samples were collected.

**Figure 2.**
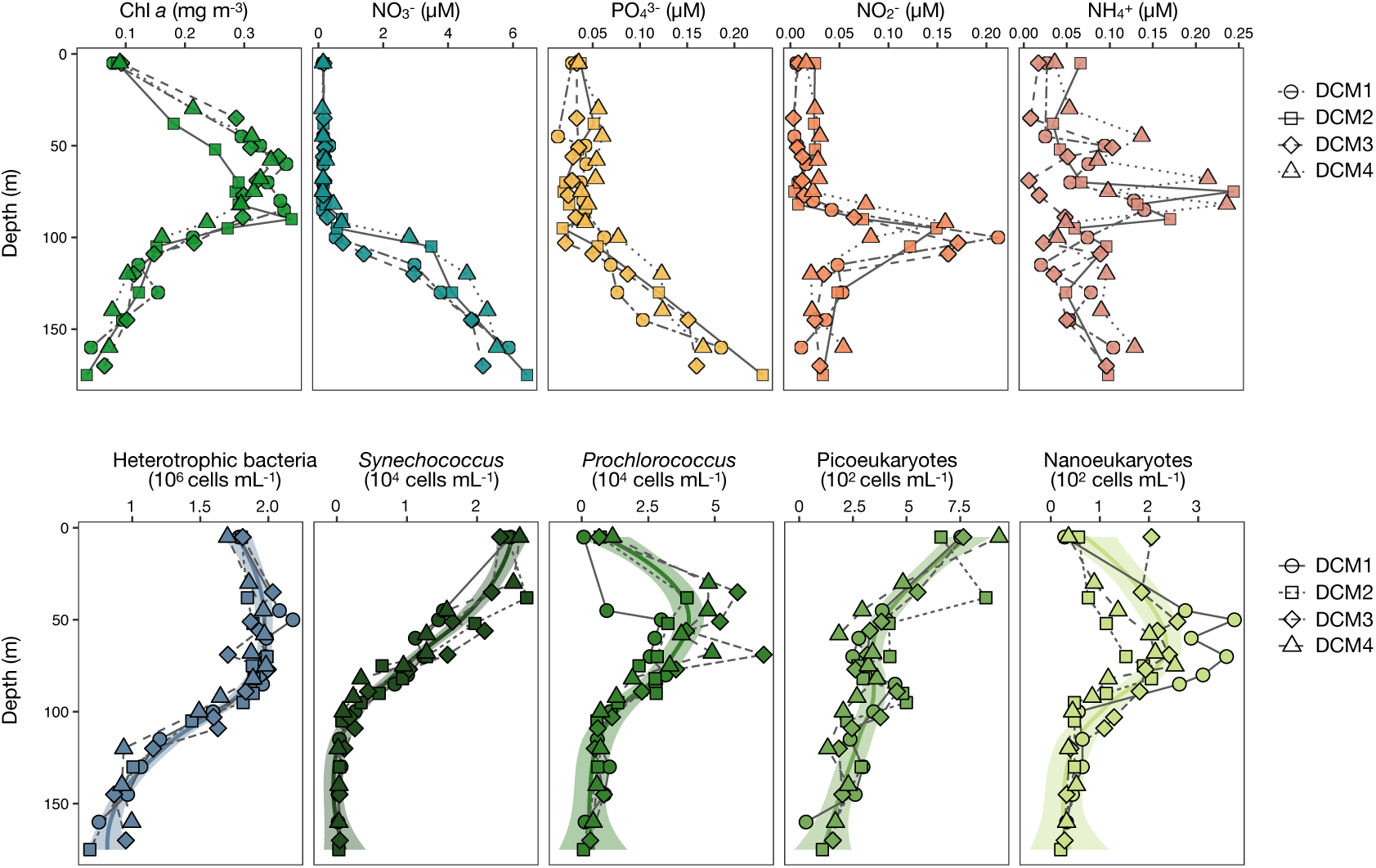
Vertical distribution of chlorophyll *a* (Chl *a*) and inorganic nutrient (nitrate, phosphate, nitrite, ammonium) concentrations (top panels) as well as microbial abundances (bottom panels) measured along the 4 vertical DCM profiles. The solid line in the lower panel represents a best fit smooth curve through the center of the data of the four DCMs calculated using weighted least squares to highlight the trends. The shaded area represents the confidence interval around the smooth.

Consistent with their broadly similar structure, nitrate and phosphate profiles showed little variation among the four DCMs (Fig. 2, Table S1). In all cases, the base of the DCM coincided with the nutricline. Nitrite concentrations remained low from the surface to just below the DCM, where they reached a maximum and then dropped abruptly (Fig. 2). This pattern was consistent across all profiles, although in DCM4 the nitrite peak occurred at a slightly shallower depth. Ammonium fluctuated throughout the water column, and exhibited the greatest variability among all measured nutrients across the four DCM profiles. Nevertheless, concentrations tended to be higher at depths coinciding with maximum values of chlorophyll *a*.

The vertical distribution of microbial abundances were quite consistent between the four DCMs, except for the abundances of photosynthetic nanoeukaryotes, which had more variability (Fig. 2, lower panel). Heterotrophic prokaryotes presented higher values in the upper 90 m (∼2·10^6^ cells mL^-1^), with the maximum values within the DCM. *Synechococcus* decreased from the surface (∼2.5·10^4^ cells mL^-1^) to the base of the DCM, where abundances became very low. In contrast, *Prochlorococcus* presented maximum abundances in the upper part of the DCMs (∼5·10^4^ cells mL^-1^, Fig. 2). Photosynthetic picoeukaryotes showed decreasing abundances from the surface (∼7.5 10^2^ cells mL^-1^) to the DCM, where they were rather constant (∼2.5·10^2^ cells mL^-1^), and then decreased slightly afterwards (Fig. 2). Although the abundance of photosynthetic nanoeukaryotes varied among profiles, they consistently showed maximum abundances (up to 2-3·10^2^ cells mL^-1^) at the depth of the DCM peak (Fig. 2).

### Changes in community structure and alpha diversity along the DCMs

The NMDS ordination of prokaryotic communities of both size fractions along the four DCMs showed a clear segregation of the samples based on size fraction and a gradual differentiation in community structure along the depth profile (Fig. 3a), supporting the view of the DCM as a microbial coenocline. In the 0.2-3 µm size fraction, the richness of prokaryotic communities showed little variation among the four DCMs, with a clear increase from the surface to 120 m depth and a slight decrease below this depth (Fig. 3b). In the 3-20 µm size fraction the richness of the prokaryotic communities was rather uniform, but showed a minimum in the 50-100 m depth layer and a slight increase with further depth (Fig. 3b). In terms of evenness, free-living (0.2-3 µm) prokaryotic communities showed higher values (i.e. less dominance) in deeper waters, whereas particle-associated (3-20 µm) communities had rather constant values throughout the DCMs structure, with a slight increase with depth (Fig. 3c).

**Figure 3.**
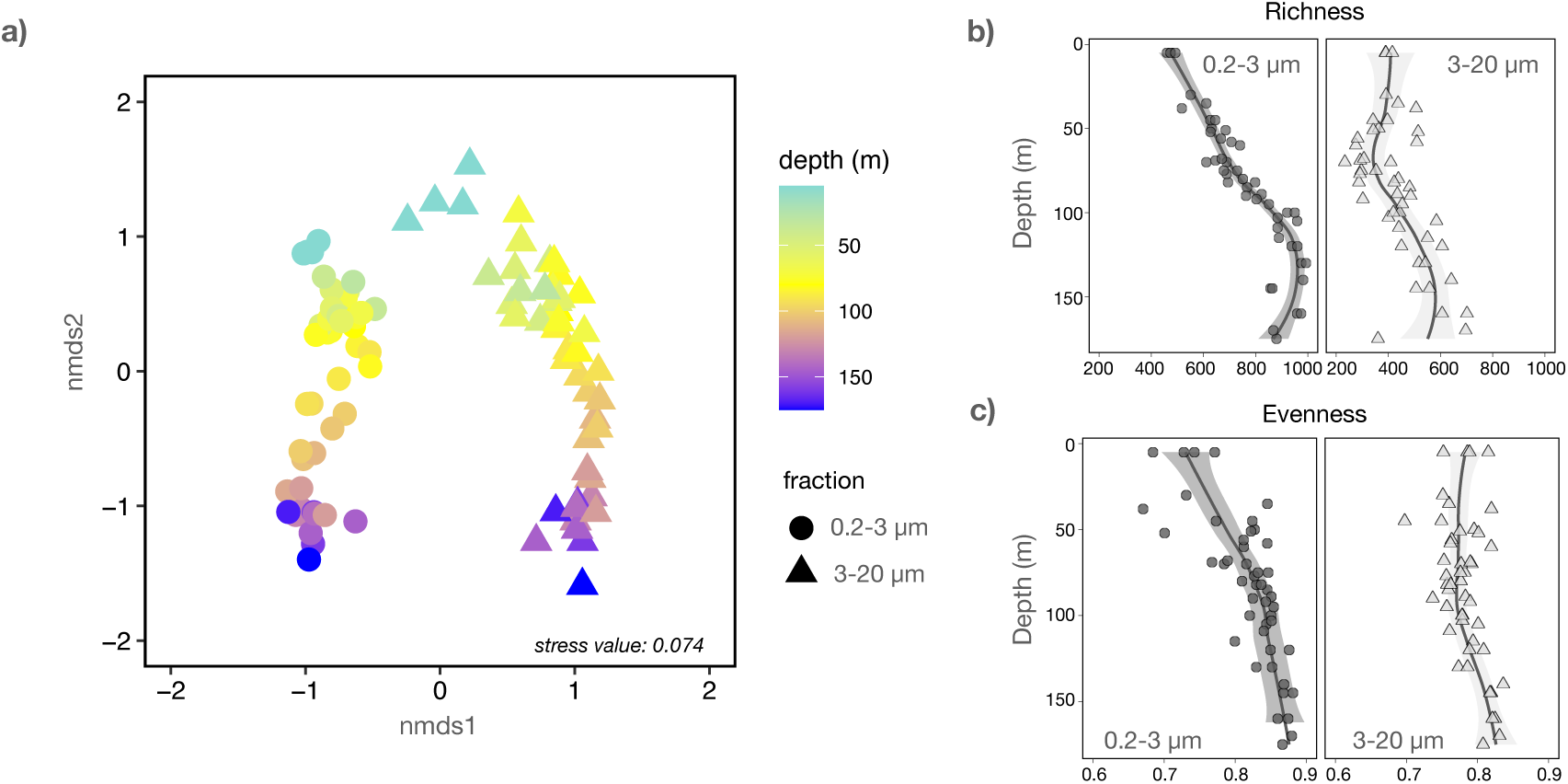
Beta and alpha diversity of the prokaryotic communities along the DCMs. **a)** NMDS ordination of the prokaryotic (16S rRNA gene based) communities in the free-living (0.2-3 µm, circles) and particle-associated (3-20 µm, triangles) size fractions. The symbols are colored based on depth. **b)** Richness and **c)** Evenness of the prokaryotic communities in both size fractions (considering the pooled four casts). Solid lines represent the best fit smooth curves through the center of the data calculated using weighted least squares to highlight the relationship between richness and evenness with depth. Grey shading represents the confidence intervals around the smooths.

### Vertical distribution patterns of prokaryotes

The taxonomic composition of the communities was quite uniform in the 4 DCMs (Fig. S3). Across both size fractions, surface waters were enriched in *Synechococcus* and Flavobacteriales, whose relative abundances generally declined with depth. In contrast, deeper layers were characterized by increased contributions of Pelagibacterales, Nitrospinota, and SAR406 in the free-living fraction, and of Planctomycetota, OM190, and Phycisphaerae in the particle-associated fraction. To explore the vertical organization at the ASV level, we grouped the prokaryotic ASV according to their vertical distribution patterns across each DCM profile using fuzzy clusters (see methods and Fig. S1 for details). This analyses revealed consistent niche segregation along the DCM gradient (Fig. 4). In the 0.2-3 µm size fraction, prokaryotic communities were partitioned into five distinct clusters, whereas four clusters were identified in the 3-20 µm size fraction. These clusters displayed similar relative contributions and vertical patterns across all four profiles, and Mantel analyses showed consistent associations with the same environmental variables across the four DCMs (Fig. S4). There was a ‘surface’ cluster that represented around 80% of the 0.2-3 µm community sequences in the surface, decreasing down to 50% of the community in the upper DCM structure, and dropping drastically below the DCM (∼100 m), where it represented less than 10% of community sequences. A surface cluster was also observed in the 3-20 µm size fraction, displaying a similar vertical distribution pattern. However, its contribution to the community at the DCM peak was lower, accounting for less than 25% of the total abundance. In both size fractions, variations in the relative contribution of this cluster to the overall community appear to be strongly driven by temperature (Fig. S4). A second cluster, labeled as ‘DCM’, comprised a small portion of the 0.2-3 µm community in the surface but peaked where chlorophyll concentrations were highest (Fig. S2), representing around 40% of the total community. Below the DCM, its relative contribution gradually decreased, mirroring the decline in chlorophyll concentrations (Fig. S2). Accordingly, variation in the distribution of this cluster was strongly associated with the chlorophyll gradient, as indicated by Mantel analyses (Fig. S4). A ‘DCM’ cluster was also identified in the 3-20 µm size fraction, where its contribution was notably higher, accounting for up to 80% of the community at its peak. A third cluster, labeled ‘below DCM’, peaked just beneath the DCM (around 100 m depth) and was generally associated to the nitrite peak (Fig. S4). This cluster accounted for over 30% of the community in the 0.2-3 µm size fraction and about 50% in the 3-20 µm size fraction. The contribution of this cluster to community sequences decreased afterwards except for the DCM2 in the 3-20 µm size fraction, where it continued increasing until 150 m before declining. A fourth cluster was formed by taxa that were rare until the base of the DCM and then started to increase in abundance (at around 80 m depth in the 0.2-3 µm size fraction and 100 m in the 3-20 µm size fraction). This cluster, designated as ‘Deep 1’, correlated with nitrate concentrations (Fig. S4) and accounted for approximately 40% of the community in the 0.2-3 µm size fraction and up to 80% of the community in the 3-20 µm size fraction. Additionally, in the 0.2-3 µm size fraction, a fifth group emerged deeper in the water column, referred to as ‘Deep 2’ (Fig. 4). The contribution of this cluster to the total community was also associated to nitrate concentrations (Fig. S4).

**Figure 4.**
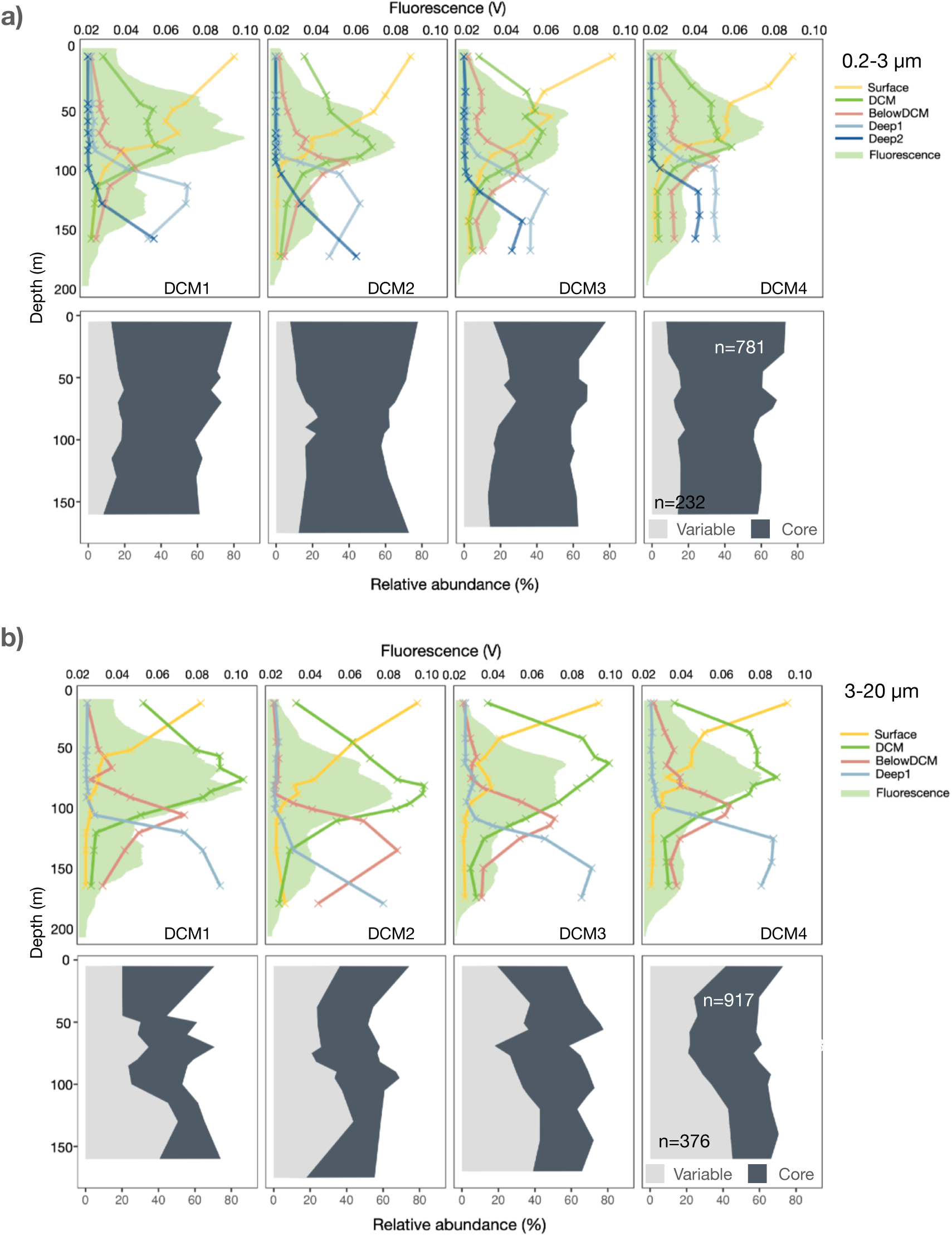
Contribution of the different fuzzy clusters to the total community across the four DCM profiles in **a)** the free-living (0.2-3 µm) fraction and **b)** the particle-associated (3-20 µm) fraction. The shaded green area represents the CTD fluorescence values at each depth. The lower panel indicates the contribution of ‘core’ ASVs (i.e., those ASVs that are consistently assigned to a single cluster across the four DCMs) and ‘variable’ ASVs (those ASVs assigned to more than one cluster). The number within the right plot indicates the number of ASVs present in each category.

### Temporal stability of vertical niches

We next investigated whether the ASVs comprising each cluster were consistently assigned to the same cluster across the four DCMs sampled, or whether their distribution shifted among clusters between the different samplings (Fig. 4, lower panels). Those ASVs that were consistently assigned to the same cluster were categorized as ‘core’, whereas those that varied were referred to as ‘variable’. In the 0.2-3 µm size fraction, we found 781 ‘core’ ASVs, that represented more than half of the total community sequences. In contrast we identified only 232 ‘variable’ ASVs, contributing approximately 20% of the community. In the 3-20 µm size fraction, 917 ASVs were categorized as ‘core’, while 376 ASVs were ‘variable’. Notably, the relative contribution of ‘core’ ASVs in this size fraction was substantially lower, particularly below the DCM.

Examining the taxonomic affiliation of the ‘core’ ASVs in the 0.2-3 µm size fraction, the ‘surface’ cluster was largely dominated by *Synechococcus*, followed by members of the Flavobacteriales, Pelagibacterales and Rhodobacterales (Fig. 5a). The taxonomic affiliation of the ‘DCM’ cluster was more diverse, with a large contribution of Flavobacteriales, followed by Verrucomicrobiota, Planctomycetes, Pseudomonadales and various orders of the Alphaproteobacteria. The ‘below DCM’ cluster was largely dominated by Pelagibacterales and Flavobacteriales, and had some representation (∼5%) of *Prochlorococcus*. The ‘Deep 1’ and ‘Deep 2’ clusters had a large contribution of the archaeal phylum Thermoproteota (formerly Crenarchaeota), which represented around 50% of the cluster in terms of sequences. The SAR324 phylum emerged in the ‘Deep 1’ and ‘Deep 2’ clusters, whereas Chloroflexota only appeared in the ‘Deep 2’ cluster. The phylum Actinomycetota was present with minor contributions in all the clusters, with highest abundances in the ‘belowDCM’ cluster. Looking at the taxonomic composition of the variable taxa, we observed that Pelagibacterales were the main contributors to this category throughout the DCM structure (Fig. 5b), followed by Flavobacteriales, *Prochlorococcus* and Pseudomonadales. SAR324 emerged in the ‘variable’ category below 120 m depth, particularly in the DCM1 and DCM2.

**Figure 5.**
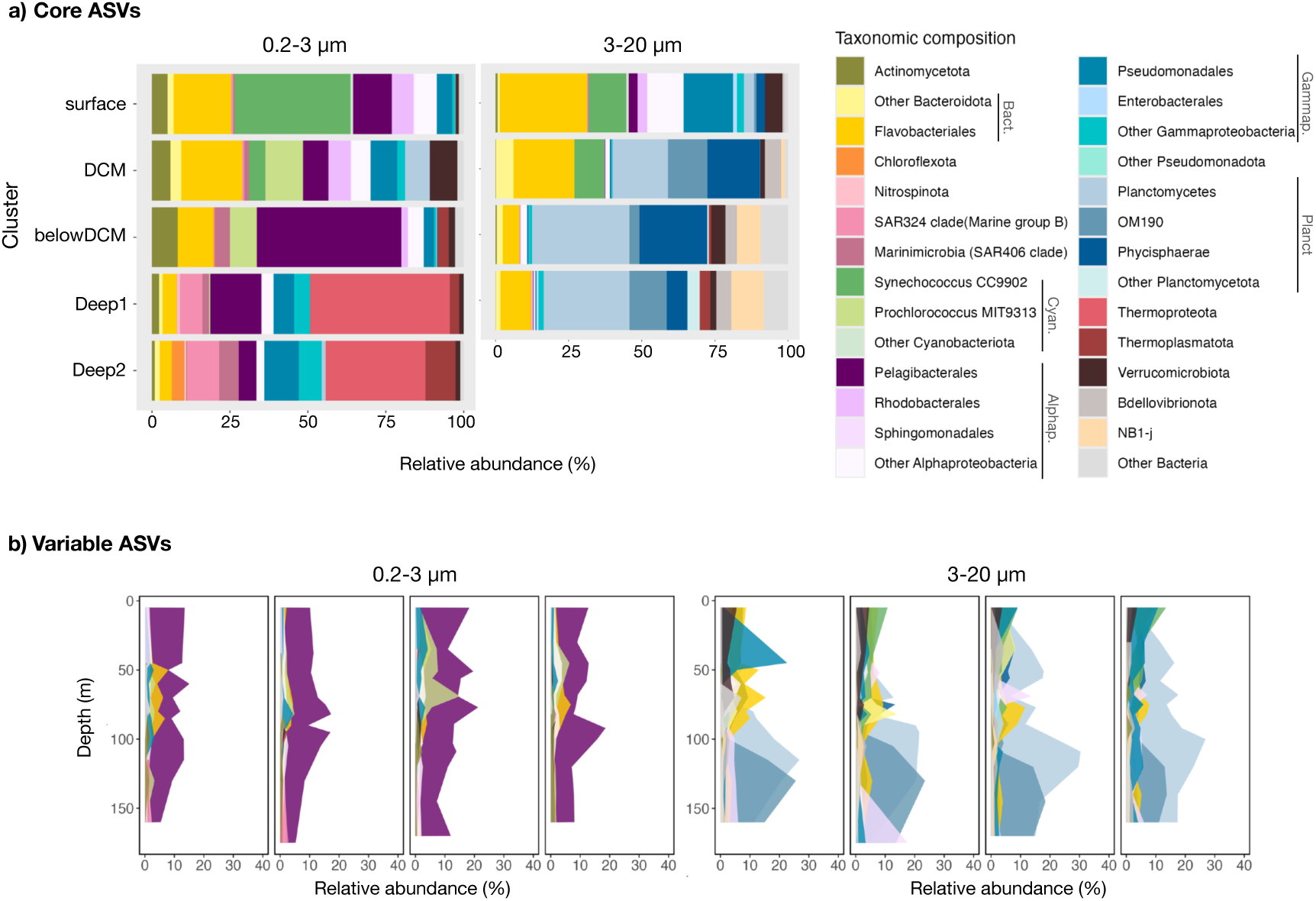
Taxonomic affiliation of ‘core’ and ‘variable’ ASVs and their contribution to prokaryotic communities in both size fractions (0.2-3 µm and 3-20 µm). **a)** Total relative abundance contributed by a given taxonomic group within each depth cluster, pooling the four DCMs. **b)** total contribution of the ‘variable’ ASVs colored by taxonomy along the different DCM profiles. Taxonomic affiliation is shown at the phylum level except for Pseudomonadota, for which both Gammaproteobacteria and Alphaproteobacteria are shown at the order level, Bacteroidota, shown at the order level, Planctomycetota, shown at the family level, and Cyanobacteriota, shown at the genus level. Taxonomic groups with low relative abundances are represented in the ‘Other Bacteria’ category.

In the 3-20 µm size fraction, the taxonomic assignation of the ‘core’ ASVs in the ‘surface’ cluster was similar to the one found in the 0.2-3 µm size fraction (Fig. 5a), except for a notably lower contribution of *Synechococcus,* a larger representation of Flavobacteriales and Pseudomonadales, and the appearance of Verrucomicrobiota. In the deeper clusters the composition drastically changed, with a large representation (over 50% of ‘core’ ASVs reads) of Planctomycetota (Phycisphaerae, OM190 and Planctomycetes). Phycisphaerae were abundant in the ‘DCM’ and ‘belowDCM’ clusters, whereas its contribution to the ‘Deep 1’ cluster was much lower. Bdellovibrionota emerged in the ‘DCM’ cluster and were also present in the deeper clusters. NB1-j were present only in the ‘belowDCM’ and ‘Deep 1’ clusters, being more abundant in the deeper one. The broad taxonomic assignation of the ‘variable’ ASV in this size fraction (Fig. 5b) was notably similar to that of the ‘core’ ASVs. In DCM1 there was a large contribution of Pseudomonadales in the upper 50 m of the water column, and this order was also prevalent along the DCM structure of DCM4. Planctomycetes and OM190 (both within the Planctomycetota) were the dominant groups, with OM190 accounting for ∼15-20% of the community sequences below 100 m depth, and Planctomycetes being present throughout most of the DCM structure. *Synechococccus* comprised part of the variable ASVs, contributing the most at the surface but being present until the base of the DCM. Sphingomonadales, practically absent in the ‘core’ category, also represented a substantial fraction of the ‘variable’ communities (Figure 5b). Verrucomicrobiota were also present, especially in the first 70 m of the DCMs. Bdellovibrionota showed a peak (up to 5% community sequences) around 30 m depth in DCM3.

Finally, we explored whether niche segregation of the individual ASVs carried a phylogenetic signal. Pairwise phylogenetic distances between ASVs increased linearly with differences in their weighted mean depth of abundance, but only across short phylogenetic distances (up to 1 arbitrary units, Fig. 6). This means that closely related ASVs tended to occupy similar depth niches, whereas more distantly related ASVs exhibit greater separation in their typical depth of occurrence. This analysis was performed using both size fractions combined, but the same pattern emerged when each size fraction was analyzed independently (Fig. S5). The observed relationship disappeared when the analyses were restricted to pairs of ASVs belonging to the same cluster (Fig. 6a). The range of phylogenetic distances over which this pattern was observed corresponded primarily to comparisons among ASVs belonging to the same order and, in some cases, the same class (Fig. 6b). Consistent with these results, ASVs assigned to the same depth cluster were generally more closely related than ASVs assigned to different clusters. For most of the dominant classes, phylogenetic distances among taxa within the same depth cluster were significantly lower than those between taxa from different clusters (Fig. S6), indicating phylogenetically conserved depth preferences. The Nitrososphaeria class was the only exception, showing the opposite pattern. This global trend was consistent when analyses were performed at the order (Fig. S7) and family (Fig. S8) levels. At the family level, however, Pelagibacterales Clade I also deviated from the general pattern, showing no significant differences between within and between cluster comparisons.

**Figure 6.**
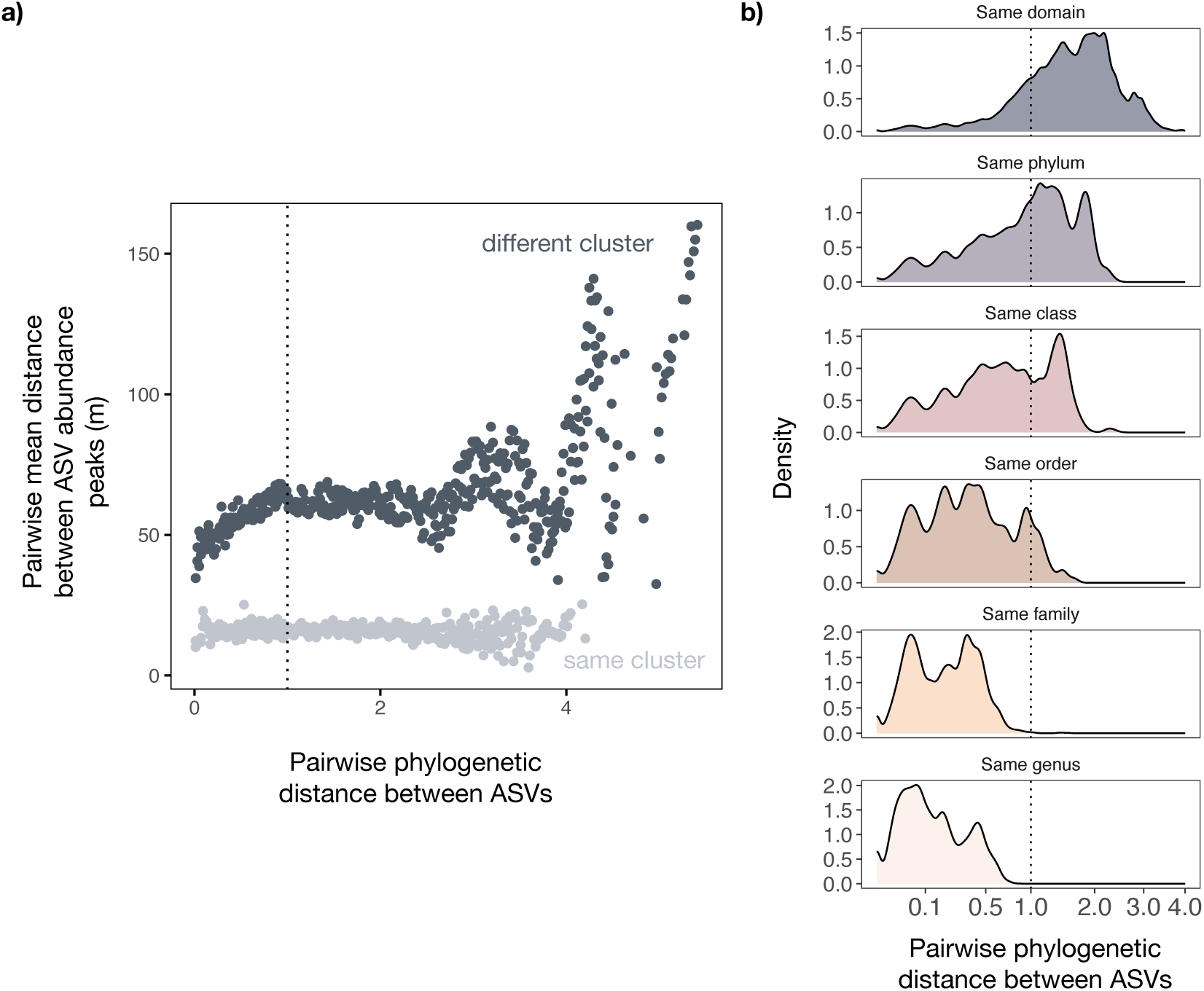
Phylogenetic signal of the vertical distribution of ASVs along the DCM (both size fractions combined). **a)** Pairwise differences in ASV depth optima, calculated as the abundance-weighted mean depth of occurrence, plotted against pairwise phylogenetic distance. Points represent mean values computed within 0.01 unit phylogenetic distance bins. Bins with less than 5 observations were discarded for the analysis. **b)** Density distributions of phylogenetic distances among ASV pairs belonging to the same Domain, Phylum, Class, Order, Family and Genus. Only the ASVs that represented more than 0.1% of relative abundance and with a cluster membership >0.75 are shown in both panels (1623 ASVs). The vertical line in panels a) and b) indicates a phylogenetic distance of 1 (until which the positive relationship between phylogenetic distance and distance in the water column is maintained).

A sharp increase in depth segregation was also apparent at phylogenetic distances above 3 units (Fig. 6a). Because this range corresponds to comparisons among very distantly related taxa (Fig. 6b), the pattern likely reflects ecological differences among major phylogenetic lineages rather than fine-scale niche differentiation among closely related ASVs.

## Discussion

Previous studies have demonstrated there is a marked vertical structuring of prokaryotic communities with depth throughout the ocean water column (e.g. Sebastián et al., 2021; Sunagawa et al., 2015). However, most research is based on samples collected from discrete layers (e.g. surface, DCM, mesopelagic and bathypelagic) or from depths separated by hundreds of meters, resulting in limited vertical resolution. This limitation is particularly relevant in the DCM, a key oceanographic feature where sharp environmental gradients, particularly in light, nutrient availability and temperature, occur over narrow depth intervals. The handful of studies that have explored the DCM at high spatial resolution have reported pronounced vertical structuring either in free-living prokaryotic communities (Haro-Moreno et al., 2018), aerobic anoxygenic phototrophic bacteria (Gazulla et al., 2026), photosynthetic picoeukaryotes (Cabello et al., 2016b) or ciliates (Dolan and Marrasé, 1995). In line with those studies, our fine-scale sampling across the DCM showed that the composition of prokaryotic communities changed progressively along this environmental gradient, in both the free-living (0.2-3 µm) and particle associated fraction (3-20 µm) (Fig. 3 and Fig. 4). This observation unveils the DCM structure as a coenocline where transitions between distinct communities occur as a continuum rather than with discrete boundaries. Community richness and evenness in the free-living (0.2-3 µm) fraction increased with depth (Fig 3b,c), consistent with previous observations reporting the same trend (Agogué et al., 2011; Sebastián et al., 2024, 2021; Severin et al., 2016; Sunagawa et al., 2015), while this pattern was not as evident in the particle-associated (3-20 µm) fraction. Particles arise from a variety of sources, including living and senescent phytoplankton cells, detritus, or marine gels (Baumas and Bizic, 2024), potentially shaping the distribution of the associated prokaryotes. However, the gradual vertical structuring observed in particle-associated communities (Fig. 3a) suggests that these communities, or the particles they inhabit, are also influenced by the environmental gradient.

The vertical changes in taxonomic composition in the free-living size fraction are in agreement with previous observations using discrete samplings (De Corte et al., 2025; Mestre et al., 2018; Sebastián et al., 2021), with a dominance of Cyanobacteria, Flavobacteriales and SAR11 in the surface and upper DCM samples, and an increase in the contribution of Thermoproteota (previously Crenarchaeota or Thaumarchaeota) below the DCM (Fig. S3). In the particle associated size fraction, communities were dominated by Planctomycetes, which are well-recognized members of the particle microbiome (Ebihara et al., 2024; Heitger et al., 2026; Mestre et al., 2018; Salazar et al., 2015) with ability to degrade complex polysaccharides (Pérez-Cruz et al., 2024; Reintjes et al., 2023).

Our fuzzy clustering approach allowed to unravel the transitions between the different communities along the DCM structure, with the possibility of studying the vertical trends at the ASV level. This approach has been applied before to analyze temporal (Gómez-Letona et al., 2025; Tobias-Hünefeldt et al., 2021) and spatial (Bagnaro et al., 2020) successions, being particularly useful at delimiting microbial transitions. ASVs grouped in different clusters along the DCM profile, linked sequentially to the temperature gradient (‘surface’ cluster), chlorophyll concentration (‘DCM’ cluster), the nitrite peak (‘belowDCM’ cluster), and the concentration of nitrate (deep clusters, Fig.4 and Fig. S4). Interestingly, in the free-living size fraction the placement of the individual ASVs along the DCM remained quite stable over the short temporal scale sampled here, with the majority of the ASVs consistently associated to the same depth cluster (‘core’ ASVs, Fig. 4 lower panel). This finding suggests that free-living prokaryotic niches remained relatively stable over the two-day period examined. Such persistence contrasts with the well documented diel variability in microbial activity within the DCM, where transcriptional activity, primary productivity and grazing pressure can change substantially over the course of a day (Becker et al., 2021; Harris, 1988; Peoples et al., 2026). In the particle-associated fraction, although most taxa exhibited niche stability (917 ‘core’ taxa vs 376 ‘variable’ taxa), the contribution of these stable taxa to the community sequences was lower than in the free-living size fraction, possibly reflecting vertical transport driven by particle sinking throughout the water column. However, this difference may also arise from the inherent higher dynamism of these communities, which usually harbor opportunistic taxa capable of rapid growth upon sudden pulses of organic matter (Leu et al., 2022).

Among taxa that did not show niche-stability, Pelagibacterales emerged as the dominant contributor in the free-living fraction, whereas Planctomycetota dominated the ‘variable’ pool of the particle associated fraction. Pelagibacterales are typical oligotrophs that have undergone genome streamlining, maintaining a minimal lifestyle focused on the efficient uptake of substrates (Grote et al., 2012), and may therefore be less influenced by the pronounced nutrient gradients along the DCM structure. Alternatively, the lack of strong niche differentiation may reflect the extensive strain-level genomic diversity within Pelagibacterales (Molina-Pardines et al., 2025), much of which is not captured by ASV-based analyses and may mask ecological specialization along the DCM gradient. Planctomycetota, on the other hand, are typically associated with large size particles (Fontanez et al., 2015), suggesting that their variable placement in the water column may result from passive transport with the particles as they sink.

The marked segregation along the DCM structure is consistent with previous results showing that, during the stratified season, most microbes in the photic zone are confined to layers approximately 30-m thick (Haro-Moreno et al., 2018). Here, however, we detected shifts in community composition over vertical scales as small as 10 m. Additionally, we showed that this niche partitioning had a phylogenetic signal (Fig. 6, Fig. S5-S8), indicating that closely related taxa tended to occupy similar habitats. This pattern in most cases extended up to the class level, and aligns with previous studies showing phylogenetic conservation in the spatial and temporal segregation of taxa in other aquatic environments (Andersson et al., 2010; Stegen et al., 2012). An exception to phylogenetic conservation was observed in Nitrososphaeria, which showed higher phylogenetic divergence among taxa occupying the same depth cluster than between different clusters. Notably, the distribution of these archaea was restricted to the two deep clusters within the 0.2-3 µm size fraction. The absence of phylogenetic conservation in the niche partitioning of this group may be linked to their evolutionary dynamics, which are strongly influenced by lateral gene transfer (LGT) followed by gene duplication (Sheridan et al., 2020). Indeed, large-scale LGT events have been proposed as major drivers of evolution across diverse archaeal lineages (Nelson-Sathi et al., 2015), including Thermoproteota (López-García et al., 2015).

## Conclusions

The DCM plays a crucial role in the structuring of the pelagic environment and consequently in the ocean carbon cycle, as it is a major site for primary production. Our fine-scale sampling approach revealed a gradual transition between distinct microbial communities across the DCM coenocline, driven by changes in environmental conditions. Closely related taxa became increasingly phylogenetically distinct with increasing separation in the water column, suggesting that water column stratification may promote fine-scale niche differentiation and potential adaptive divergence along the vertical gradient. Investigating these short spatial scales is therefore essential for understanding microbial niche partitioning in the ocean and its potential implications for biogeochemical processes. Since ocean warming may alter the structure of the DCM, particularly the depth at which it forms (Richon et al., 2019), this could potentially lead to shifts in microbial community composition and, consequently, ecosystem functioning.

## Acknowledgements

We thank everybody onboard the *R/V* García del Cid for their help. Special thanks go to the ship crew and the UTM technicians for smooth operation of the CTD. We also extend our gratitude to the MARBITS bioinformatics platform at the Institut de Ciències del Mar.

## Author contributions

IF, OS and JMG designed the sampling. CC, CMV and IF collected the samples. CMV and VB performed the nucleic acid extraction. CMV did the DADA2 processing. MS conceived the idea of the manuscript and together with CMV and AO performed the analyses. MS did the figures and wrote the manuscript. All authors have revised and approved the final version.

## Data availability

Sequences are available at ENA under accession number PRJEB122873 and metadata associated to these sequences are in table S1.

