## Supplementary information for "Fine-scale niche partitioning of prokaryotic communities across the deep chlorophyll maximum coenocline"

This file contains:

Supplementary methods
Eight supplementary figures

**Supplementary methods:**

*Chlorophyll a and inorganic nutrient measurements*

For chlorophyll *a* (Chl a) measurements, 200 mL of water were filtered through a Whatman GF/F filter and stored frozen on board until analysis. The pigment was extracted in cold acetone (90% v/v) for 24 h and analyzed with a 10 AU Turner Designs bench fluorometer, previously calibrated with pure Chl *a* (Sigma Aldrich) (Holm-Hansen et al., 1965). Samples for nitrate, nitrite, ammonium, phosphate and silicate were collected in 10 mL polyethylene bottles and kept at -20°C until analysis. Their concentrations were determined using a CFA Bran + Luebbe autoanalyzer following the methods described by (Hansen and Koroleff (2007).

*Heterotrophic bacteria and picophytoplankton abundance*

For the determination of heterotrophic bacteria and picophytoplankton abundance, samples (1.8 mL) were fixed using a solution of 1% paraformaldehyde + 0.05% glutaraldehyde (final concentration), deep frozen in liquid nitrogen and stored at −80 °C until analyzed. Subsamples for heterotrophic bacteria were stained with SybrGreen I (Molecular Probes, final concentration 1000× dilution of the commercial product) for some minutes in the dark. For picophytoplankton analyses, subsamples were unstained. Cell counting was performed by using a BD FACSCalibur cytometer with a 488 nm laser emission as detailed in Gasol and Morán (2015).

*DNA extraction, 16S rRNA gene amplification and sequencing*

Water samples (2 L) were sequentially filtered with a peristaltic pump through a 20 µm mesh onto 3 µm and then 0.2 µm polycarbonate filters, which were then kept frozen in liquid nitrogen on board and stored at -80ºC once in the laboratory. DNA was extracted with the phenol-chloroform protocol following Massana et al., (1997). The 515F-Y and 926R primers (Parada et al., 2016) were used to generate amplicons targeting the V4-V5 region of the 16S rRNA gene and sequenced in a Illumina Miseq platform at RLT Genomics (Lubbock, TX, USA; http://rtlgenomics.com/). Primers and spurious sequences were trimmed using *cutadapt* (Martin, 2011). To differentiate exact sequence variants DADA2 v1.8 was used (Callahan et al., 2016) which resolves ASVs by modelling the errors in Illumina-sequenced amplicon reads. Taxonomic assignment was performed using the *assignTaxonomy* function against SILVA v.138.2 (Quast et al., 2012).

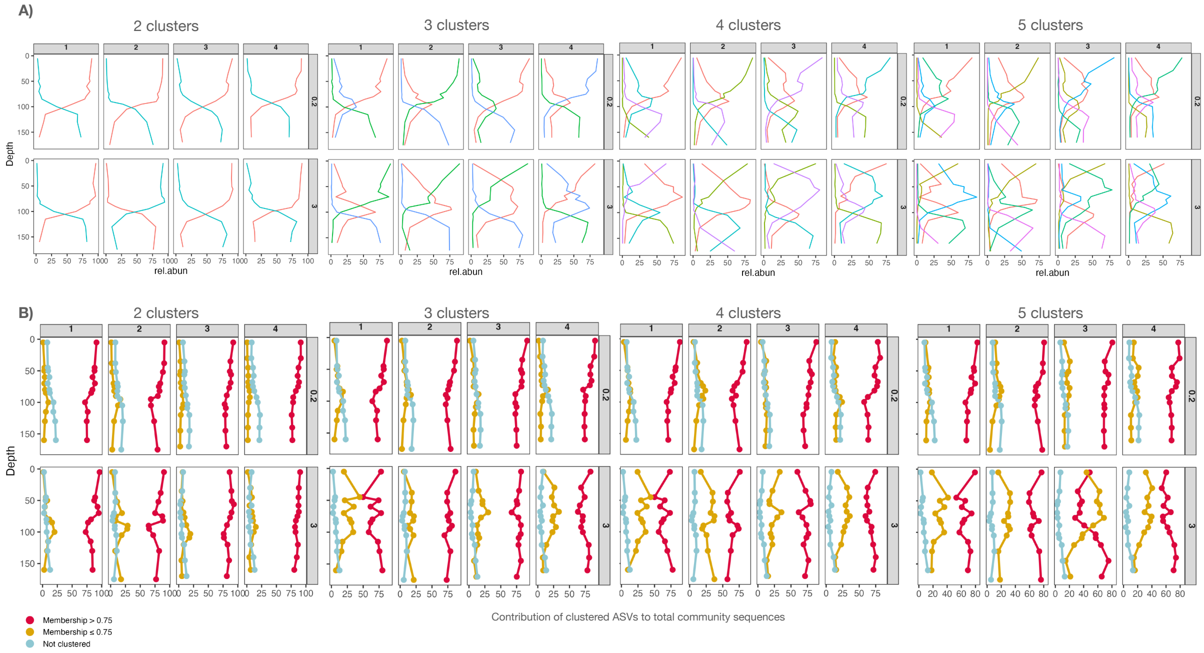


**Figure S1.** Procedure for the selection of the appropriate number of clusters. A) Grouped abundances of the ASVs in 2, 3, 4 and 5 clusters for the 0.2-3 µm size fraction (upper panel) and the 3-20 µm size fraction (lower panel). B) the proportion of ASVs with a cluster membership score >0.75 (red), <0.75 (yellow) or not clustered (light blue). To choose the number of clusters we looked for distinct non-redundant trends, while maintaining a high proportion of ASVs with a high membership score. 5 clusters were chosen for the 0.2-3 µm size fraction, because this number allowed for the detection of the belowDCM cluster, which contained a population that represented up to 30% of the community, while the membership score was maintained high throughout the DCM structure. For the 3-20 µm size fraction 4 clusters were chosen, as the trend was noisier and the membership score decreased notably, particularly in DCM3 if we chose 5 clusters.

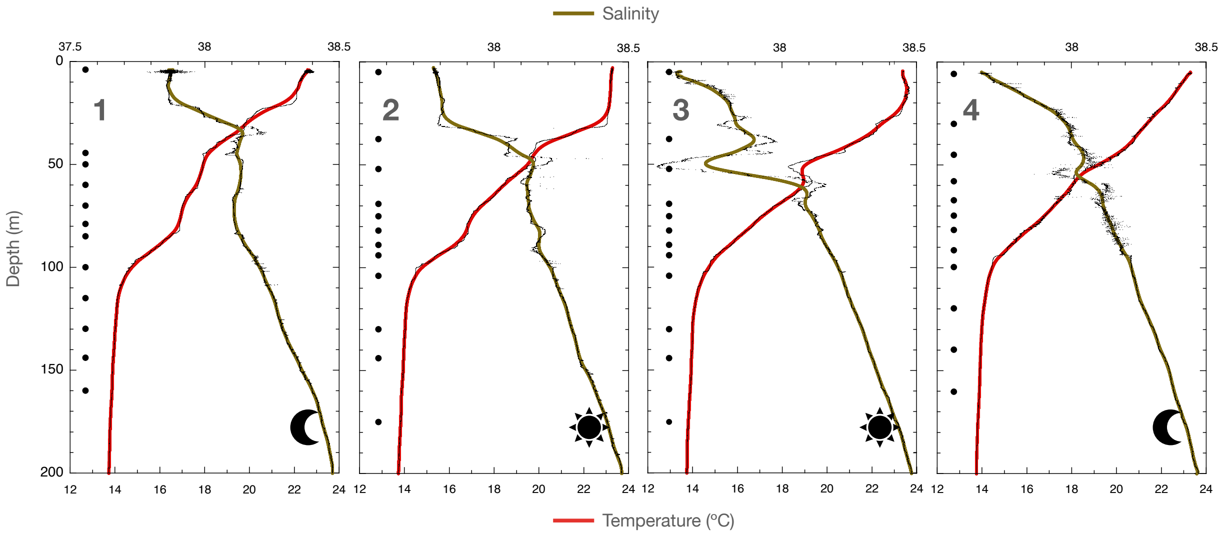


**Figure S2**. Temperature and salinity profiles of the 4 DCM casts. The number in each panel indicates the DCM number. Black dots indicate the depths at which seawater samples were collected. The lines correspond to smoothed Stineman interpolations combined with a local geometric weight covering 10% of the data range using the original CTD data (seen in the background) computed by the software Kaleidagraph (vs. 5.01, Synergy Software. (2026). https://www.synergy.com/)

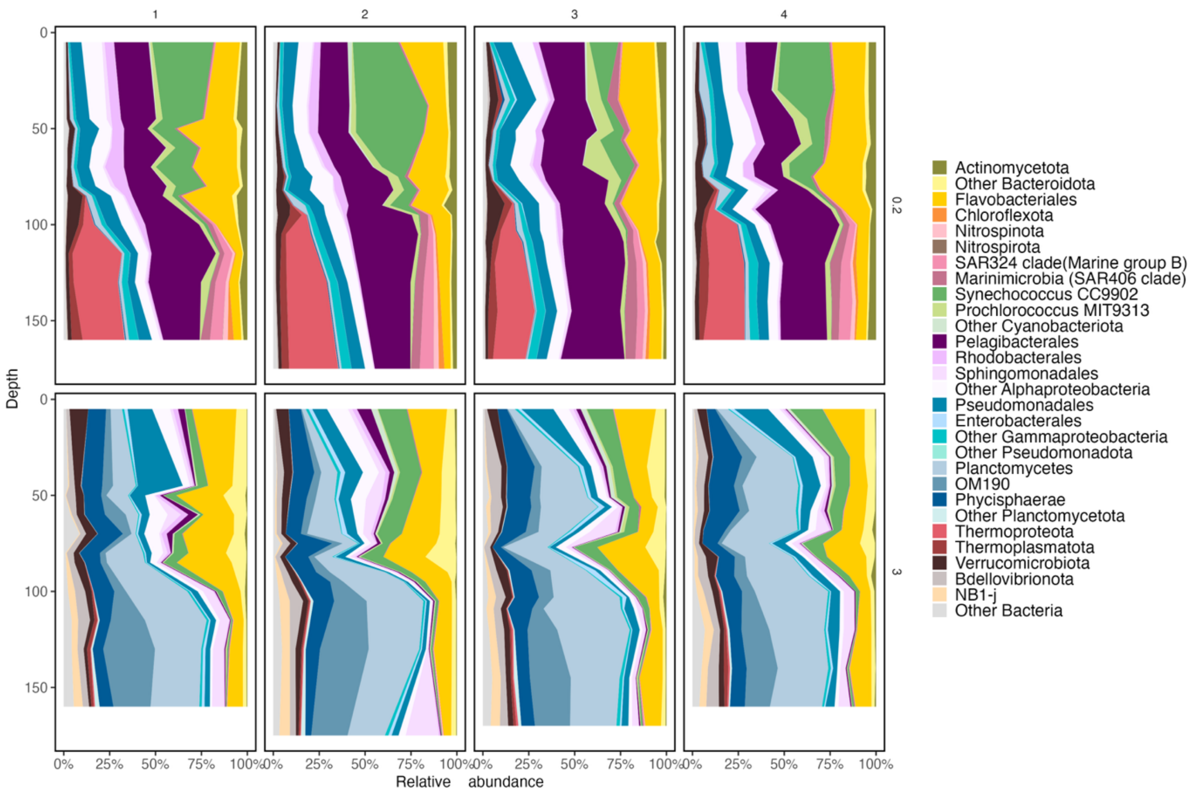


**Figure S3**. Taxonomic composition of the prokaryotic communities across the four DCMs in the 0.2-3 µm size fraction (upper panels) and the 3-20 µm size fraction (lower panels). Taxonomic affiliation is shown at the phylum level except for Pseudomonadota, for which both Gammaproteobacteria and Alphaproteobacteria are shown at the order level, Bacteroidota, shown at the order level, Planctomycetota, shown at the family level, and Cyanobacteriota, shown at the genus level. Taxonomic groups with low relative abundances are represented in the ‘Other Bacteria’ category.

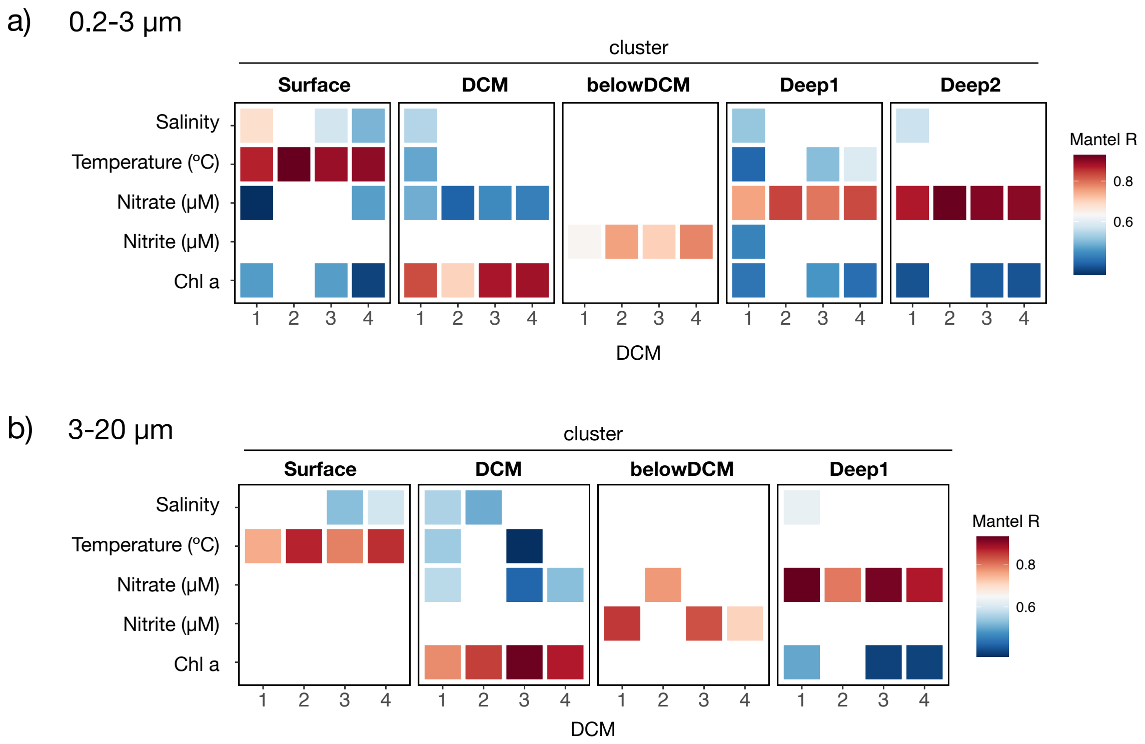


**Figure S4.** Role of environmental variables and chlorophyll *a* (Chl a) in shaping the cluster distribution across the DCM structure. Heatmap showing R coefficients of the Mantel correlations between the Euclidean distances of the summed contribution of the ASVs belonging to each of the clusters and the Euclidean distances of the environmental variables measured across samples in each individually sampled DCM. Only significant correlations are shown (Bonferroni-adjusted $p < 0.05$).

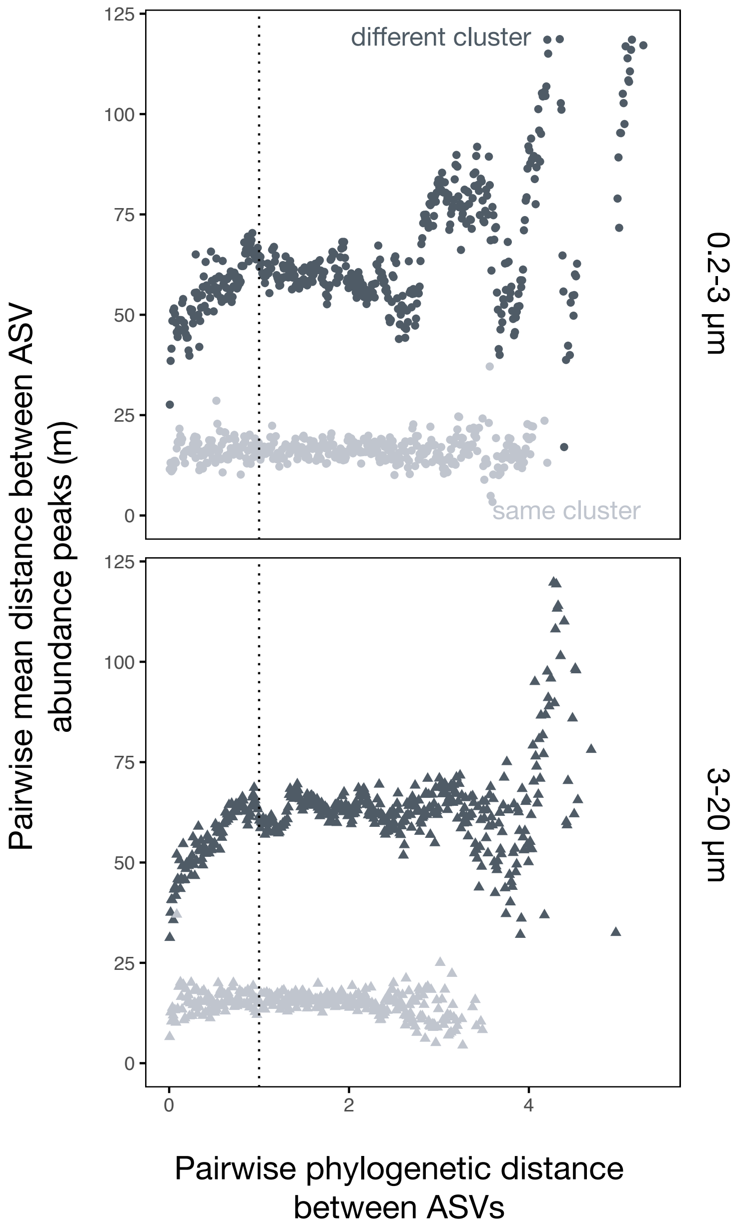


**Figure S5**. Phylogenetic signal of the vertical distribution of ASVs along the DCM separated into the free-living (0.2-3 µm) and particle-associated (3-20 µm) size fractions. Pairwise differences in ASV depth optima, calculated as the abundance-weighted mean depth of occurrence, plotted againsts pairwise phylogenetic distance. Points represent mean values computed within 0.01 unit phylogenetic distance bins (only bins with more than 5 observations were considered). Only the ASVs that represented more than 0.1% of relative abundance and with a cluster membership >0.75 are shown in both panels (1623 ASVs). The dotted vertical line indicates a phylogenetic distance of 1 (up to which the positive relationship between phylogenetic distance and distance in the water column is maintained).

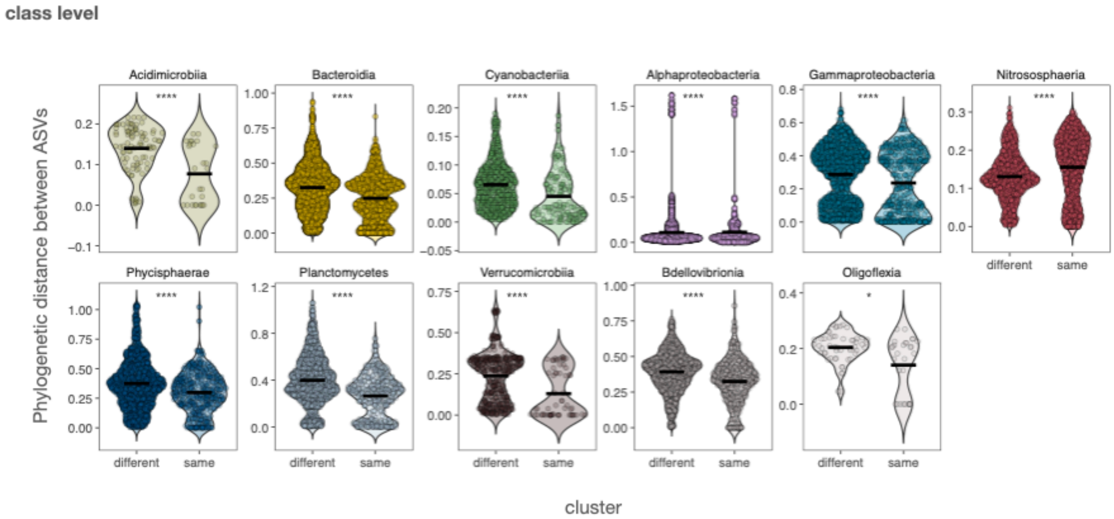


**Figure S6.** Phylogenetic distances between ASVs belonging to the same depth cluster or different clusters grouped at the class level. The analyses was performed including only families with more than 20 comparisons in each cluster are shown. Asterisks denote Benjamini-Hochberg (BH) p-adjusted significance of the Wilcoxon tests (*: <0.05, **: <0.01, ***: <0.001, ****: <0.0001, ns: p>0.05). Violin-plots are colored based on the taxonomic affiliation (same colors as in the main text). The cross-bar indicates the mean.

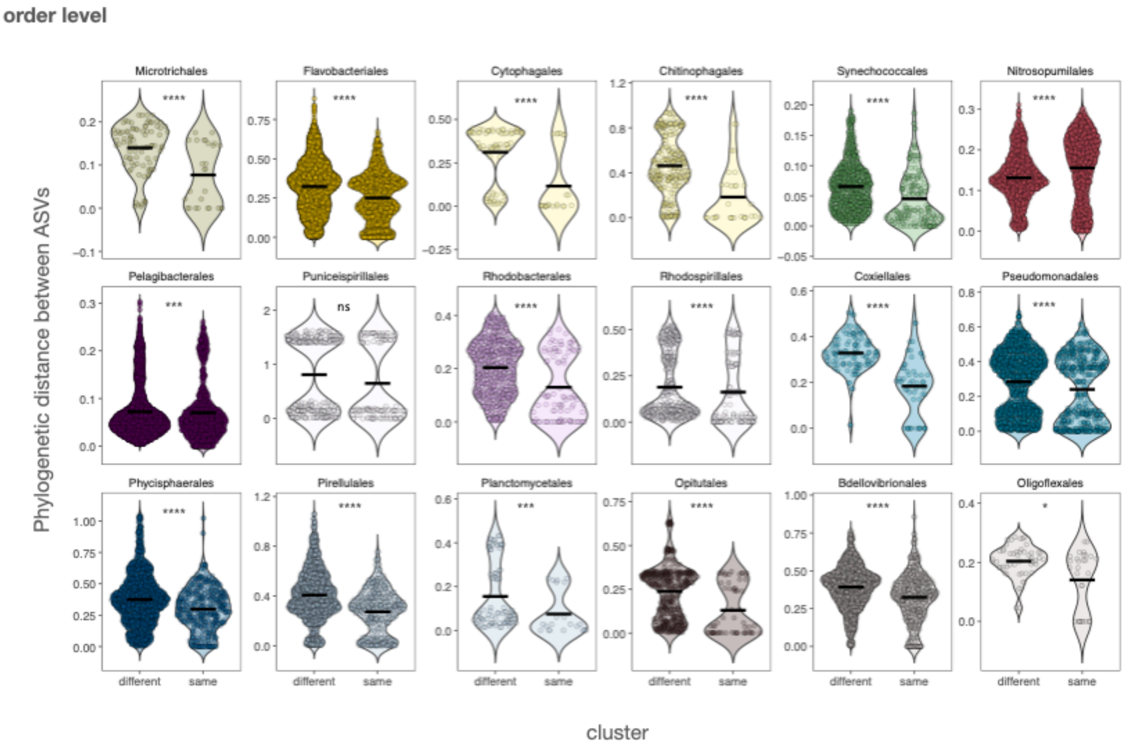


**Figure S7.** Phylogenetic distances between ASVs belonging to the same depth cluster or different clusters grouped at the order level. The analyses was performed including only families with more than 20 comparisons in each cluster are shown. Asterisks denote Benjamini-Hochberg (BH) p-adjusted significance of the Wilcoxon tests (*: <0.05, **: <0.01, ***: <0.001, ****: <0.0001, ns: p>0.05). Violin-plots are colored based on the taxonomic affiliation (same colors as in the main text). The cross-bar indicates the mean.

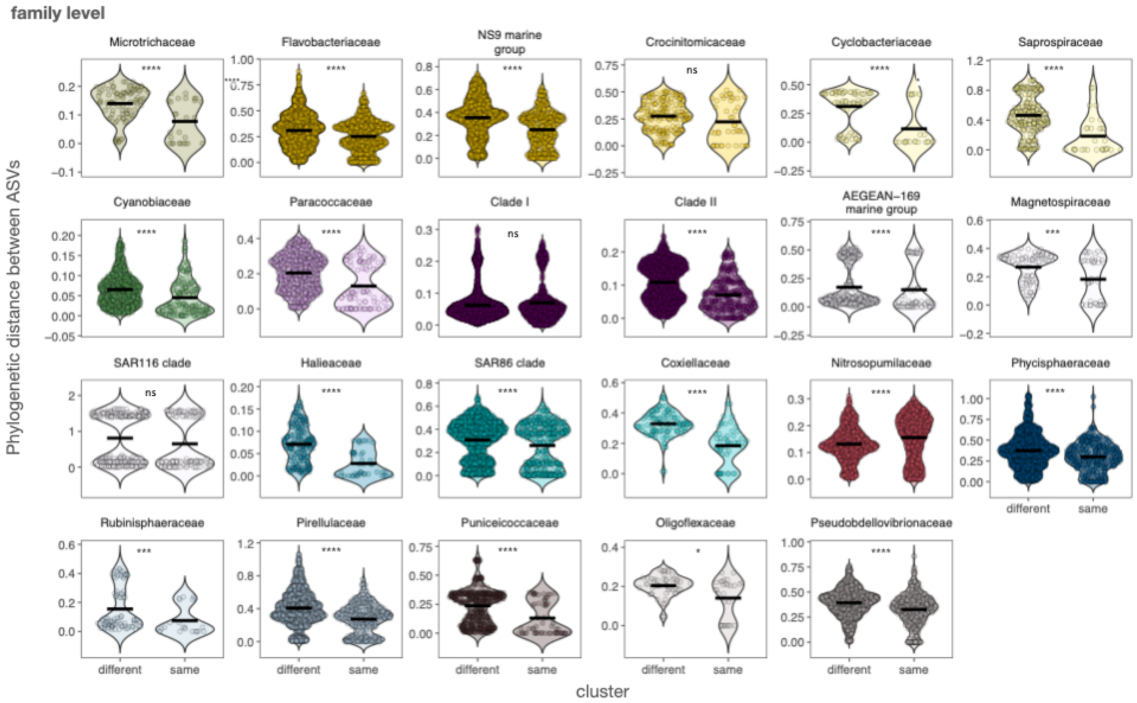


**Figure S8.** Phylogenetic distances between ASVs belonging to the same depth cluster or different clusters grouped at the family level. Only families with more than 20 comparisons in each cluster are shown. Asterisks denote Benjamini-Hochberg (BH) p-adjusted significance of the Wilcoxon tests (*: <0.05, **: <0.01, ***: <0.001, ****: <0.0001, ns: p>0.05). Violin-plots are colored based on the taxonomic affiliation (same colors as in the main text). The cross-bar indicates the mean.
